# Foliar mRNA spray induces protein synthesis in monocot crop and dicot model plant species

**DOI:** 10.1101/2025.08.07.668951

**Authors:** Veli Vural Uslu, Moritz Dobrowitsch, Kathrin Preussel Danger, Alexandra Ursula Furch, Jonas Noetzold, Antje Maria Richter, Timo Schlemmer, Yihan Dong, Gabi Krczal, Aline Koch

**Author notes:** equal contribution (Veli Vural Uslu, Moritz Dobrowitsch, Kathrin Preussel Danger). **Corresponding Author(s):** Aline Koch, Veli Vural Uslu.

## Abstract

*In planta* gene expression via exogenous mRNAs has a wide range of potential biotechnological applications from genome editing to plant protection and to circumventing tissue culture. Yet, regarding mRNA delivery into intact plant cells, the purity, stability and translational efficiency of exogenous mRNAs appear as obstacles for effective exogenous mRNA application. Our study investigates the potential of exogenous mRNAs in mediating protein synthesis in plant cells. We applied unformulated, unmodified green fluorescent protein (GFP)-coding mRNA to monocot (*Triticum aestivum, Hordeum vulgare, Zea mays*) crops and dicot model plant *Arabidopsis thaliana* and observed GFP signals in all species, confirming the successful uptake and translation of mRNA into protein. This approach opens possibilities for broader, GM-free applications in plant science.

## Introduction

Plants are surprisingly receptive to foreign RNA molecules. The application of double-stranded RNA (dsRNA) and small interfering RNA (siRNA) foliar sprays has demonstrated that these exogenous RNAs can cross plant barriers and trigger RNA interference (RNAi), silencing genes in pests and pathogens (Koch et al., 2016; Rank and Koch 2021, Uslu et al. 2025). This breakthrough has led to the development of the first commercial RNA spray-based plant protection products, such as Calantha™, which is now authorised in several markets (Yan et al. 2024; Narva et al. 2025). These developments offer a non-GM alternative to synthetic pesticides and highlight the ability of plants to absorb nucleic acids from external sources.

Based on our observations that plants absorb sprayed RNAs largely unspecifically and passively, we hypothesized that this phenomenon could extend beyond non-coding RNAs. Could coding RNAs, such as messenger RNAs (mRNAs), also enter plant cells and be translated into functional proteins? This concept could transiently restore gene functions, produce protective proteins and accelerate breeding or functional studies without stable transformation, which is critical where genetic engineering faces scientific, economic and societal limitations (Qaim 2020; Lassoued et al. 2021).

Here, we demonstrate that unmodified, unformulated mRNA, when applied as a simple leaf spray, can enter plant cells and be translated into protein. Using an mRNA encoding green fluorescent protein (GFP), we treated three monocot crop species, wheat (*Triticum aestivum*), barley (*Hordeum vulgare*) and maize (*Zea mays*) and the model plant species *Arabidopsis thaliana*. Remarkably, the translation of GFP mRNA into GFP protein was observed in all the plants tested under confocal laser scanning microscopy (CLSM), western blot, and/or ribosome profiling, indicating efficient uptake, stability and cytoplasmic translation.

To our knowledge, this is the first demonstration of foliar-applied naked mRNA being internalized and expressed in plant cells. Unlike previous reports which required encapsulation or formulation for delivery, our findings reveal the inherent ability of plants to internalize and utilize foreign RNA species, including coding mRNA. These results broaden the conceptual framework of RNA-based technologies in agriculture and pave the way for diverse, GM-free applications in basic research and crop improvement.

## Results

### In-vitro transcribed mRNA constructs can be translated in plant extract and in protoplasts

For foliar spray applications, we produced both capped and uncapped synthetic mRNAs *in vitro* that contained only the coding sequence (CDS) of the respective fluorescent reporter protein (eGFP or citrine), followed by a C-terminal triple hemagglutinin (HA) tag to enable immunodetection (Fig. 1a). To confirm the compatibility of these synthetic mRNA molecules with plant translation machinery, translation assays were conducted both *in vitro* using wheat germ extract (WGE) and *in vivo* using barley protoplasts. In the in vitro assay, protein translation was confirmed by immunoblotting with an anti-HA antibody (Fig. 1b). For the in vivo analysis, protoplasts were isolated from barley leaves and treated with citrine mRNA via passive co-incubation or PEG 4000-mediated transfection in combination with a non-coding carrier plasmid. Fifteen hours after treatment, expression of the encoded protein was detected using confocal laser scanning microscopy (CLSM) (Fig. 1c).

**Figure 1:**
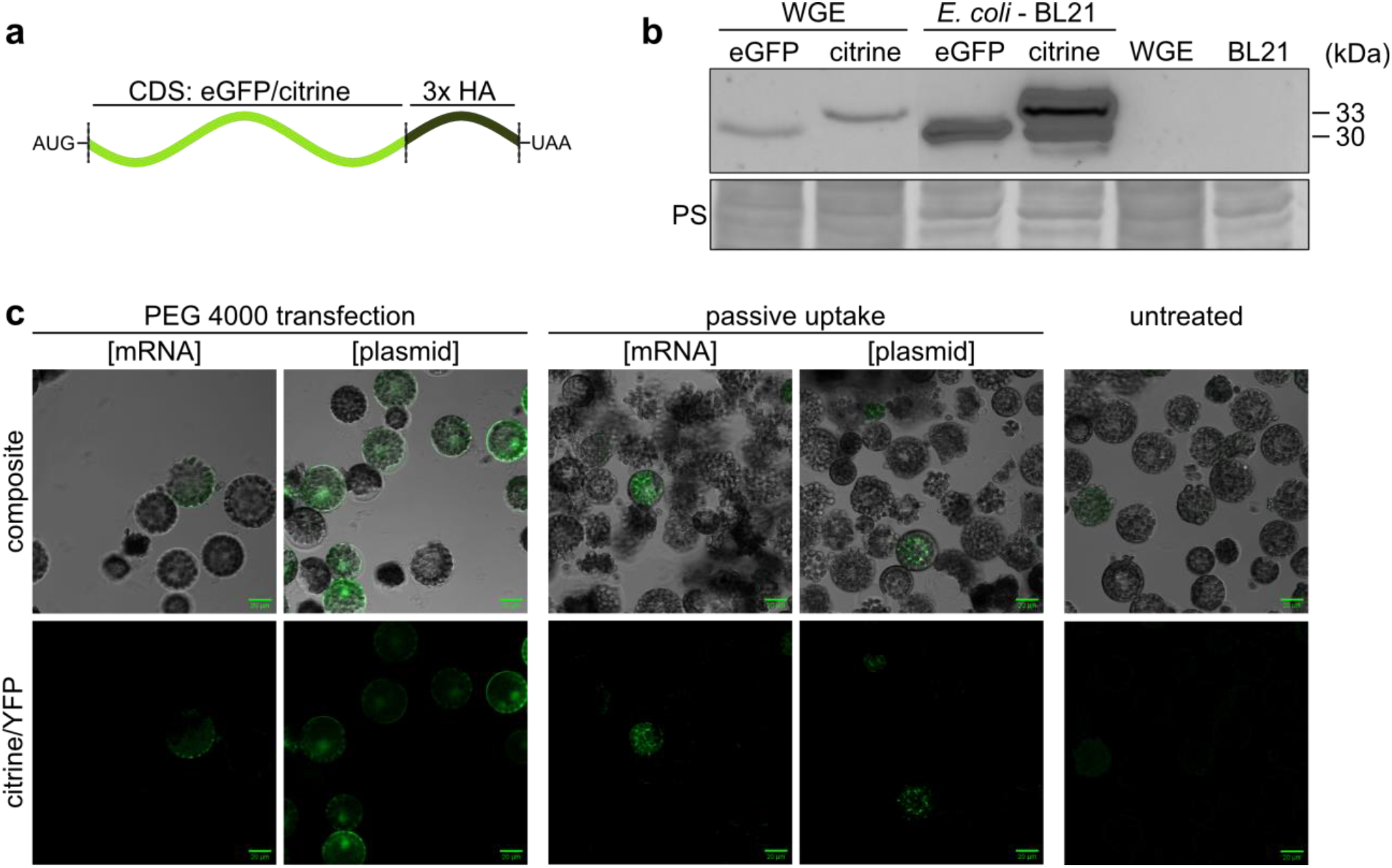
*In vitro* transcribed mRNAs are functional in cell-free and cellular systems. (**a**) Schematic representation of synthetic mRNA constructs encoding eGFP or citrine, each carrying a C-terminal 3×HA tag. (**b**) Western blot showing *in vitro* translation products from 5 µg mRNA in wheat germ extract. Controls: Total protein from *E. coli* BL21 cells overexpressing the respective proteins (100 µg). Detection was performed with primary anti-HA (mouse) and secondary anti-mouse peroxidase (goat) antibodies. Expected molecular weights: eGFP – 30.2 kDa; citrine – 32.7 kDa. Ponceau S staining confirms equal loading. (**c**) Barley protoplasts from 2-week-old seedlings were treated with citrine mRNA via PEG-mediated transfection (with carrier plasmid) or passive incubation. A constitutively citrine-expressing plasmid served as a positive control. CLSM images were taken 15 h post-treatment. Citrine fluorescence was excited at 514 nm (2.6% YFP laser intensity), detected at 508–588 nm. Scale bar: 20 µm.

### GFP signals detected in vascular and epidermal leaf tissues following mRNA spray application in barley, wheat and maize

In order to evaluate the potential for externally applied mRNA to be translated in plants, we initially sprayed in vitro transcribed green fluorescent protein (GFP) mRNA (717 nucleotides) onto the leaves of barley (*Hordeum vulgare*), wheat (*Triticum aestivum*) and maize (*Zea mays*). Previous studies have shown that barley can efficiently absorb sprayed RNAs, such as dsRNA and siRNA, even without the use of formulations or surfactants (Koch et al., 2016, 2019; Höfle et al., 2020; Biedenkopf et al., 2020; Werner et al., 2020). In line with these findings, we detected GFP-positive cells in barley leaves, whether or not the commercial surfactant Silwet was present (see Fig. 3A–C). This confirms that unmodified mRNA can also enter cells via passive uptake. We have previously demonstrated that non-coding RNAs enter plant tissues via the stomata and diffuse into the apoplast. From there, Dicer-derived siRNAs indicated intracellular processing and an apoplast-to-symplast transition. Consistent with these mechanisms, we observed a GFP signal in epidermal (Fig. 2A), subepidermal and vascular tissues, suggesting that coding mRNAs may also be taken up in a similar manner. Interestingly, GFP signals were also detected in vascular bundles and phloem-associated cells, indicating the presence of mRNA or its translated product in inner tissues (Fig. 3B, C).

**Figure 2:**
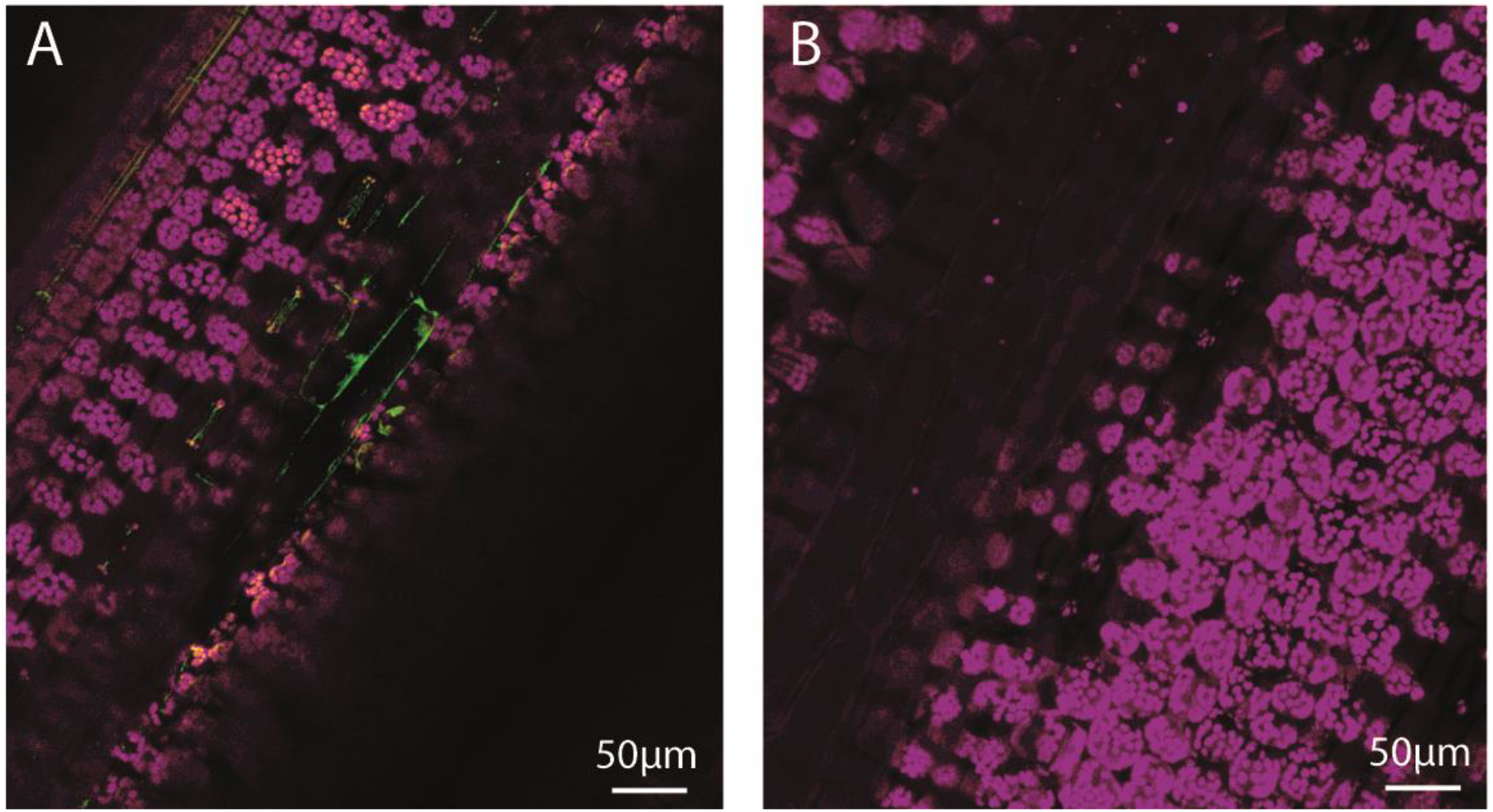
Confocal laser scanning microscopy (CLSM) reveals the uptake of exogenously applied GFP-mRNA into the epidermal cells of barley leaves. The adaxial site of barley leaves were sprayed with *in vitro* transcribed GFP mRNA (717 nucleotides) and analyzed for GFP fluorescence 24 hours after treatment. (A) GFP-positive signals (green) are visible in the adaxial epidermal and stomata cells and cavities. Specificity of the observed signals was confirmed in lambda scanning mode. (B) The mock treated control shows no GFP signal. Scale bars 50 µm.

**Figure 3:**
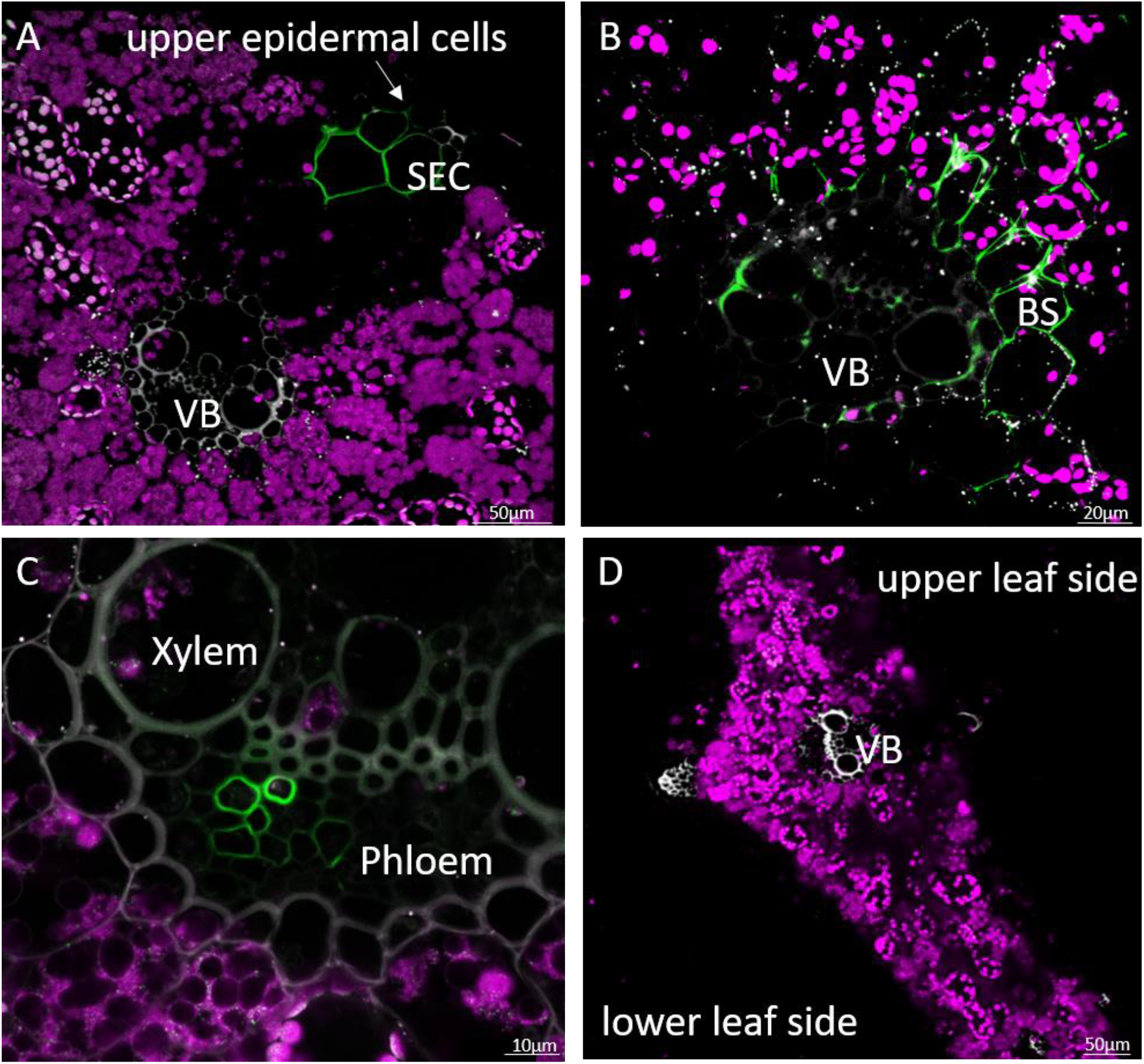
Confocal laser scanning microscopy (CLSM) reveals the uptake of exogenously applied GFP-mRNA and its subsequent distribution within barley leaves. The leaves were sprayed with *in vitro* transcribed GFP mRNA (717 nucleotides) and analyzed for GFP fluorescence 24 hours after treatment. Cross-sections of the leaves were analyzed using CLSM. (A) GFP-positive signals (green) are visible in subepidermal cells (SEC) and in the upper epidermis above the vascular bundles (VB), indicating effective uptake without the need for surfactants. (B) Signal detection in bundle sheath (BS) cells and vascular tissue suggests that the mRNA is accessible to inner tissues. (C) The presence of GFP fluorescence in phloem cells near the xylem implies the potential movement of mRNA or translated protein towards vascular tissues. (D) The mock-treated control shows no GFP signal, confirming the specificity of the observed fluorescence in the treated samples. Scale bars: A = 50 µm; B = 10 µm; C = 10 µm; D = 50 µm

However, since the GFP protein itself is known to move between cells, further work is required to distinguish between the movement of RNA and its resulting protein. To determine whether this observation also applies to wheat, a species generally considered to be less permeable to sprayed RNAs, we performed similar foliar mRNA applications. CLSM analyses of cross-sections revealed GFP fluorescence in wheat leaves 24 hours after treatment (Fig. 4A–C). The GFP signal was detected in subepidermal and mesophyll cells, as well as in cells surrounding the vascular bundles, including the phloem and xylem. Signals were also found in stomatal cavities and adjacent epidermal cells, suggesting that entry through stomata may facilitate mRNA uptake in wheat as well as in barley.

**Figure 4:**
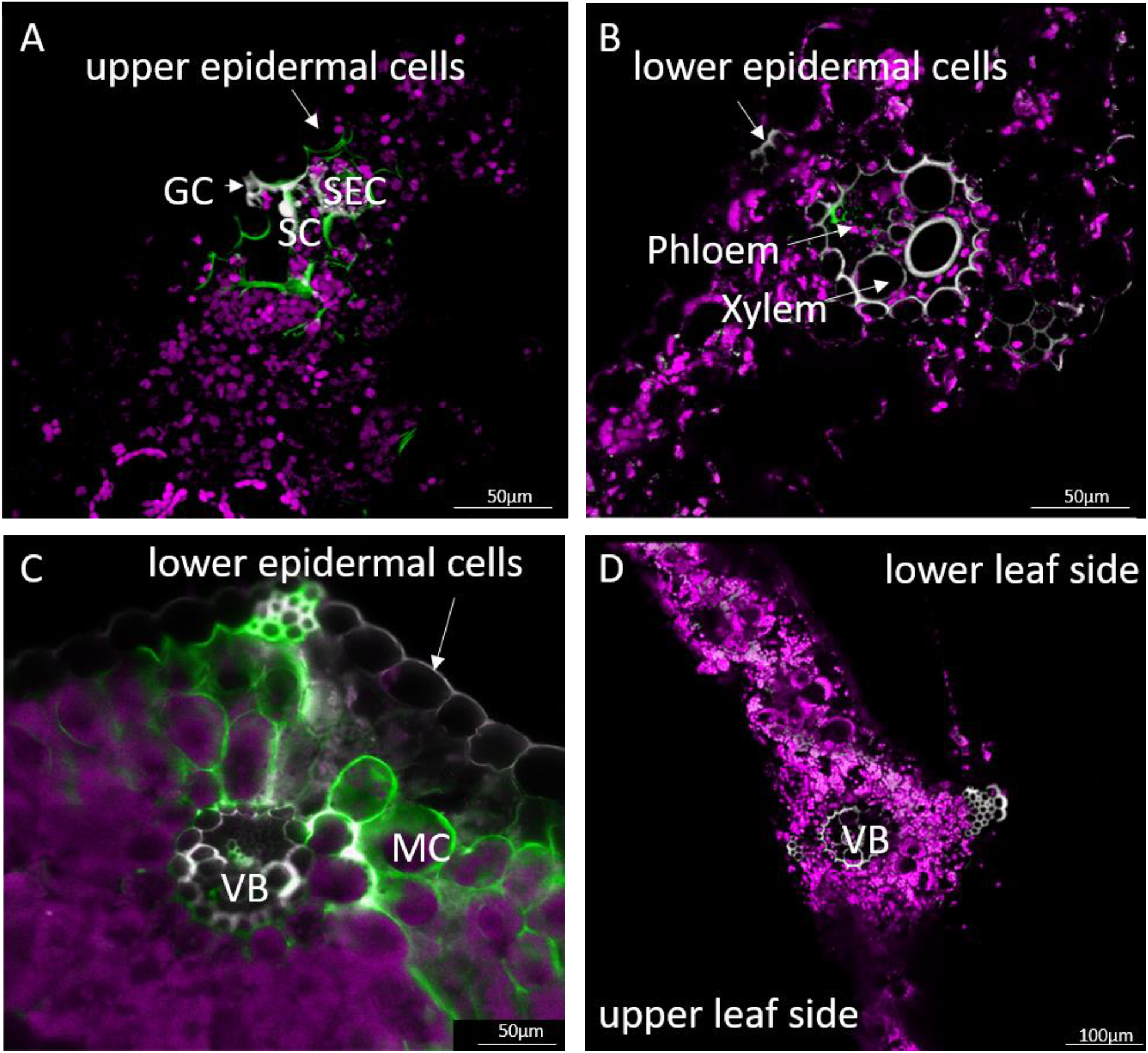
Detection of GFP-positive cells after foliar mRNA spraying on wheat leaves. Confocal laser scanning microscopy (CLSM) images show cross-sections of wheat leaves 24 hours after the foliar application of *in vitro* transcribed GFP mRNA. The green fluorescence indicates the presence of GFP (A) shows the GFP signal detected in subepidermal cells (SEC) and stomatal cavities (SC) beneath the upper epidermis. GC = guard cells. (B) GFP fluorescence is observed in vascular tissues, including the phloem and xylem, as well as in adjacent lower epidermal cells. (C) The GFP signal is visible in mesophyll cells (MC), subepidermal cells and cells surrounding the vascular bundle (VB) in the lower leaf area. (D) Mock-treated control showing no GFP signal. Scale bars A-C = 50 µm; D = 100 µm.

Subsequently, the foliar spray application of mRNA on the monocot maize was conducted to further verify the putative ubiquitous feasibility of mRNA spray applications. CLSM analyses of cross sections in maize leaves 24 hours after treatment revealed GFP fluorescence in the vascular bundles, including the phloem and xylem (Fig. 5 A-B). These results suggest that barley, wheat and maize leaves can absorb and distribute externally applied coding RNAs despite their anatomical and physiological differences. Taken together, these findings confirm that exogenously applied mRNA can be taken up and expressed in these three major cereal crops, suggesting that this strategy could be applicable to many monocot species.

**Figure 5:**
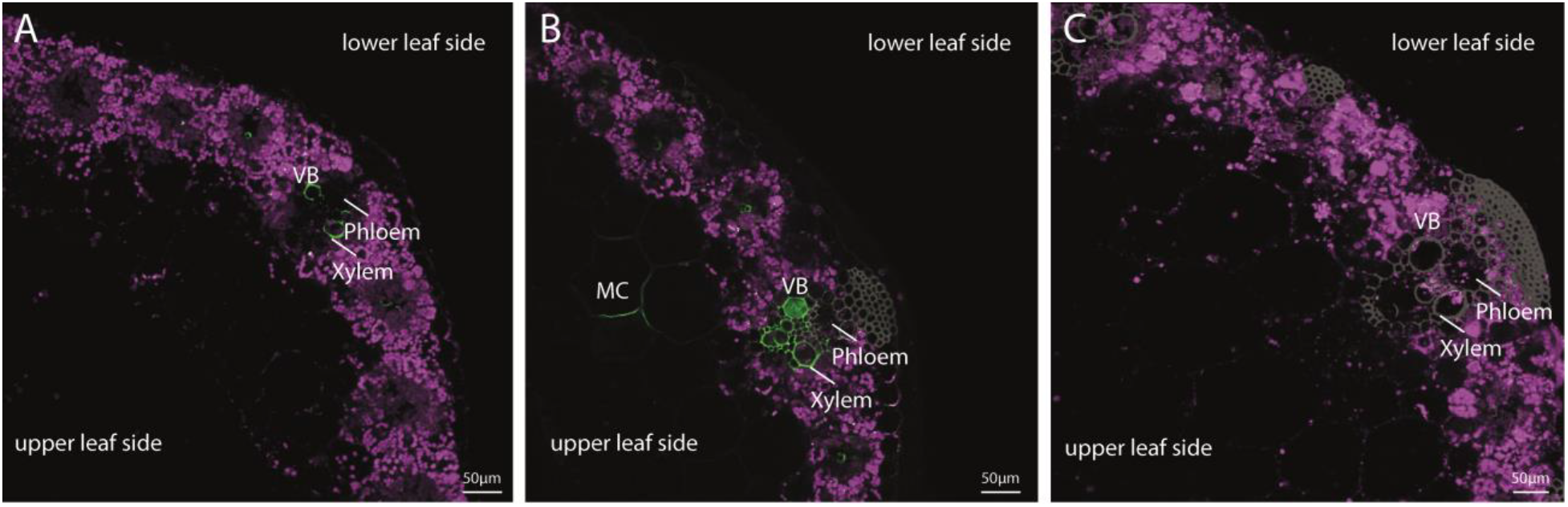
Detection of GFP-positive cells after foliar mRNA spraying on maize leaves. Confocal laser scanning microscopy (CLSM) images show cross-sections of maize leaves 24 hours after the foliar application of *in-vitro* transcribed GFP mRNA. The green fluorescence indicates the presence of GFP signal (A) 24 hours and (B) 48 hours in the vascular tissues, including the phloem and xylem. (C) Mock treated control showing no GFP signal. (VB) vascular bundle; (MC) mesophyll cells. Scale bars 50 µm.

### Ribosome profiling of Arabidopsis leaves following spraying with GFP mRNA revealed enhanced translation in *rdr6* mutants

As a dicot model, we delivered GFP mRNA to Arabidopsis rosette leaves using high-pressure spraying (Uslu et al. 2022). However, due to the mechanical damage caused by spraying, leaves treated with water alone exhibited elevated background fluorescence of GFP under imaging, which complicated direct detection of GFP. Therefore, we used ribosome profiling to quantitatively analyze the uptake and translation of GFP mRNA 48 hours after treatment (Fig. 6). We used sucrose gradient centrifugation to separate ribosomal complexes into heavy polysomes (HP), light polysomes (LP), monosomes (80S) and ribosomal subunits (60S and 40S), as well as a supernatant fraction (TOP). RNA was extracted from each fraction (n = 3 biological replicates per condition), reverse-transcribed, and transcript abundance was quantified by RT-qPCR for GFP and the housekeeping genes Act2 and Ubq10. Transcript levels in each fraction are expressed as a percentage of the total cellular mRNA (the sum of all fractions). In water-treated controls, GFP transcripts were undetectable across all ribosomal fractions, whereas Act2 and Ubq10 showed canonical distributions, predominantly in heavy and light polysomes, indicating active translation. When GFP mRNA was sprayed onto Col-0 leaves, the distributions of the housekeeping genes remained unchanged and GFP transcripts were mainly detected in the light polysome fractions, indicating translation initiation but limited polysome loading. Treatment with puromycin, a translation inhibitor that releases ribosomes from mRNA, shifted GFP transcripts from polysomal fractions towards monosomal and supernatant fractions, thus validating their ribosome association and active translation status. As RDR6 mediates post-transcriptional gene silencing via RNA-dependent RNA polymerase and subsequent RNA interference (RNAi), we hypothesized that RDR6 reduces the pool of translationally active sprayed GFP mRNA by converting it into double-stranded RNA precursors for degradation. Consistent with this hypothesis, the *rdr6-11* mutant exhibited a significant increase in GFP mRNA occupancy within heavy polysomes compared to Col-0 (p < 0.05, Student’s t-test), indicative of enhanced translation efficiency. Furthermore, Act2 and Ubq10 transcripts in rdr6-11 also shifted towards heavier polysomal fractions compared to the wild type, suggesting that RDR6 deficiency has a broad influence on mRNA translation dynamics.

**Figure 6:**
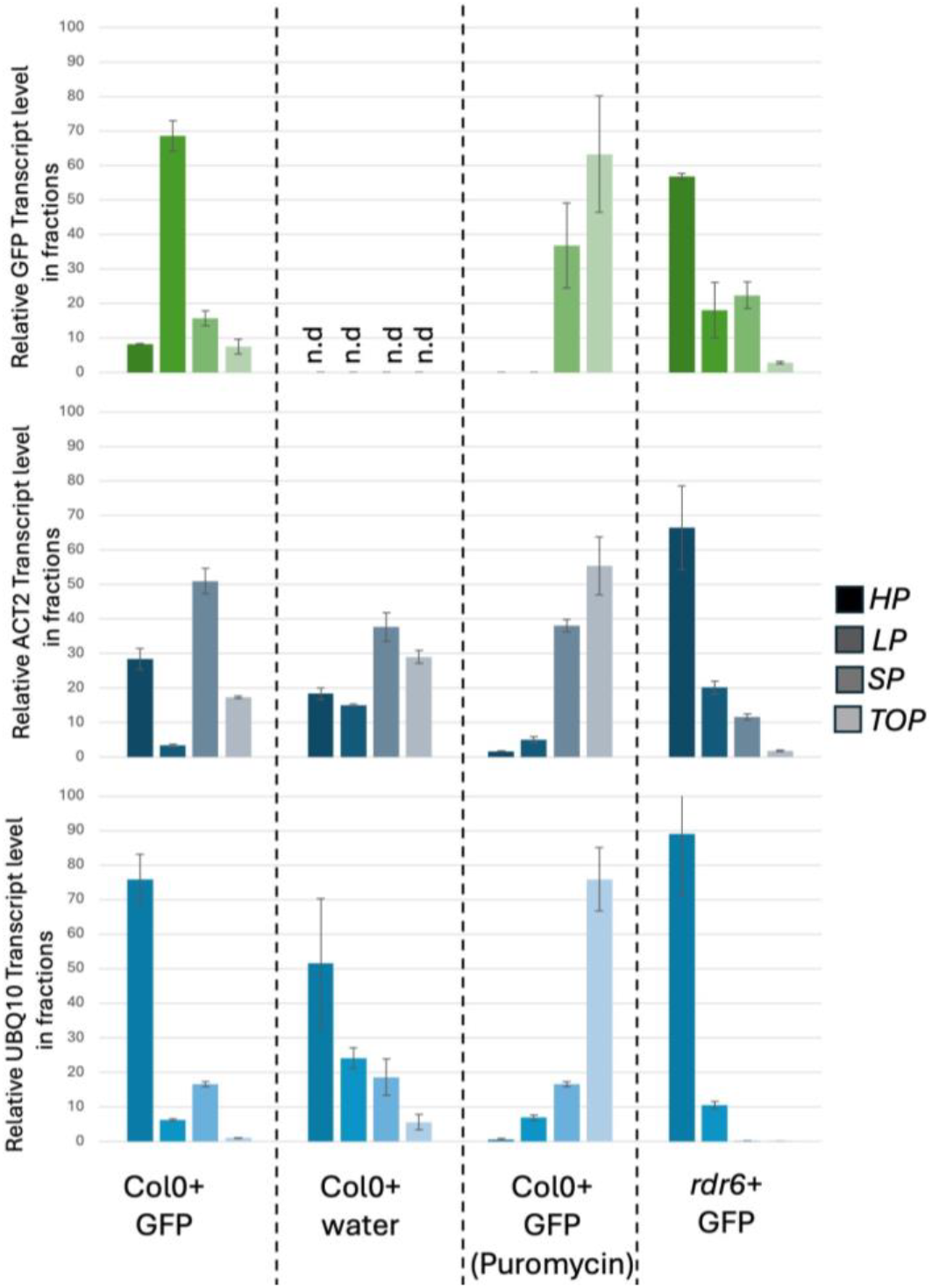
Ribosome Profiling of *Arabidopsis thaliana* rosette leaves upon spraying of GFP mRNA. GFP mRNA or water control was delivered to *A*.*thaliana* rosette leaves by high pressure spraying technique (HPST). 48h post treatment, three different ribosomal fractions and one supernatant fraction was collected in a sucrose gradient: Heavy Polysomes (HP), Light Polysomes (LP), 80S, 60S and 40S ribosomal fragments (SP), Supernatant (TOP). Color intensities for green, gray and blue follow the depicted sample legend in gray. In each sample (N=3 for each condition), total mRNA level was set to 100 (HP+LP+SP+Top). RNA extracts from each fragment were reverse-transcribed and quantitative PCR was used to assess the level of occupancy of GFP (green) and two housekeeping genes, ACT2 (dark blue) and UBQ10 (blue). in each fraction, the value shows a percentage in total mRNA level. Puromycin treatment dissociated polysomes mRNAs are released to upper fractions. Col0 + GFP refers to HPST of GFP mRNA on Col0, Col0+ water refers to HPST of water on Col0, Puromycin treatment is used to verify that mRNAs. rdr6-11 (Col0 background) is a homozygous deletion of RNA-Dependent-RNA polymerase 6. Error bars indicate Standard error.

Taken together, these results demonstrate that exogenously applied GFP mRNA is taken up and translated in Arabidopsis leaves and that RDR6 acts as a negative regulator of translation by channeling foreign mRNA into RNAi pathways, thereby limiting polysome loading and translation efficiency.

## Discussion

Our findings show that unmodified, unformulated mRNA can enter plant cells and be translated into functional proteins when applied as a simple foliar spray. This fundamentally challenges the perception that plant cells are highly selective against the uptake of foreign RNA. Instead, our data suggest that there is a more permissive system for RNA uptake, opening the door to the application of multiple RNA classes beyond mRNA. These include non-coding RNAs, such as circular antisense RNAs (caRNAs) (Hossain et al., 2025), which, due to their covalently closed structure, may be more stable within plants and the environment. This would enhance the persistence and efficacy of RNA-based treatments. Previous studies have demonstrated that plants can absorb long double-stranded RNAs (dsRNAs) measuring around 1.3 kb (Höfle et al., 2020). This lends weight to the idea of fusing protein-coding sequences to dsRNAs, paving the way for multifunctional RNA sprays. This could deliver multiple functional RNAs simultaneously, including components for CRISPR/Cas9-mediated gene editing, regulatory RNAs, and RNA-binding proteins (RBPs) that stabilize therapeutic dsRNAs within plants and potentially across kingdoms.

The broad transferability of our approach is demonstrated by the effective uptake and translation of mRNA in both monocot (e.g. barley, wheat and maize) and dicot (*Arabidopsis thaliana*) species. This suggests that RNA spray applications could be widely applicable across diverse crops, overcoming the species-specific limitations of transgenic technologies. The potential applications of sprayable mRNA extend well beyond plant protection and stress resilience. These include transient protein expression to enhance nutritional value (e.g. vitamins or allergen-free variants), delivery of genome-editing components (e.g. CRISPR/Cas systems) for transient trait engineering without genomic integration, and expression of RBPs to stabilize co-delivered dsRNAs targeting pathogens or pests. Furthermore, the possibility of cross-kingdom delivery of plant-expressed proteins could allow interactions with symbionts, pathogens or herbivores to be manipulated. Spray-based delivery of mRNA provides a unique opportunity to study protein-protein interactions and test heterologous protein function in a cross-species context, eliminating the need for stable transformation. For example, transient translation of protein complexes in leaves could enable rapid screening of functional domains or synthetic biology constructs. Compared to Agrobacterium-mediated transient expression (agroinfiltration), direct mRNA spraying bypasses host range restrictions and biosafety concerns associated with bacterial delivery systems, making it a more attractive option for field applications.

Despite these promising findings, several fundamental questions remain unanswered and must be addressed to realize the full potential of RNA spray technologies. Most critically, the precise mechanisms by which exogenous RNA molecules enter plant cells remain unclear. Our observation that exogenously applied mRNA is selectively degraded in rdr6-overexpressing lines suggests that components of the RNAi machinery may play an active role in modulating RNA stability and possibly uptake. However, the cellular pathways and plant surface structures involved in RNA internalization remain unknown. Similarly, the mechanisms underlying cell-to-cell or systemic transport of exogenous RNA and the extent to which sprayed RNA can move within plant tissues are poorly understood. Recent studies by Borniego et al. (2024) have shown that endogenous RNAs, including tRNAs, mRNAs and siRNAs, are actively secreted onto the leaf surface in a non-vesicular manner to form RNA condensates that may interact with surface microbiota. Furthermore, distinct subpopulations of extracellular vesicles (EVs) appear to be implicated in plant immune signaling, yet they do not co-transport small RNAs (Koch et al. 2025). This raises questions about whether sprayed RNAs interact with comparable extracellular compartments. The extent to which these natural RNA secretion and transport pathways affect the fate, mobility, or biological activity of exogenously applied RNA remains to be determined. Understanding such interactions could provide valuable insights into optimizing RNA formulations and delivery strategies in both basic and applied plant science, including crop improvement, breeding, and microbiome engineering.

To realize the full potential of mRNA spray technology, future efforts must also focus on improving RNA stability, translational efficiency, and tissue-specific targeting. Protective chemical modifications and innovative formulation approaches, such as lipid nanoparticles or biodegradable polymers, will be essential for enhancing environmental robustness and achieving effective translation *in vivo*. Together, these innovations could transform RNA sprays into a powerful, field-deployable platform for precision agriculture and plant biotechnology.

## Materials and Methods

### Plant material and growth conditions

Second leaves of 2-week-old barley (*Hordeum vulgare*, cultivar Golden Promise) and wheat (*Triticum aestivum* L.) were detached and transferred to square Petri plates (120 × 120 × 17 mm) containing 0.7% water agar. Barley plants were grown under controlled conditions with a 16 h light photoperiod (240 µmol m^−2^ s^−1^ photon flux density), day/night temperatures of 18 °C/14 °C, and 65% relative humidity.

### mRNA synthesis

StemMACS™ eGFP mRNA (Miltenyi Biotec) was diluted to 100 ng/µl in H_2_O according to the manufacturer’s instructions. For comparison, GFP mRNA was also generated in-house from a plasmid backbone. GFP coding sequences were amplified by high-fidelity Phusion PCR (ThermoFisher) using primers designed with Benchling (primer3-based, www.benchling.com). The forward primers contained the T7 promoter sequence (TAATACGACTCACTATAGG), and the reverse primers were specific for GFP from the pEGFP vector (Clontech; https://www.addgene.org/vector-database/2488/). PCR conditions followed ThermoFisher’s Phusion DNA polymerase annealing recommendations (www.thermofisher.com/tmcalculator). PCR products were size-verified by agarose gel electrophoresis, purified by phenol-chloroform extraction followed by ethanol precipitation. Plasmid templates for in vitro transcription included a T7 promoter upstream of the reporter coding sequence fused to a triple hemagglutinin (3xHA) tag (pT7-eGFP-3xHA-tag, pT7-citrine-3xHA-tag). Plasmids were linearized downstream of the insert using restriction enzymes to enable T7 run-off transcription. In vitro transcription (IVT) was performed either with the mMESSAGE mMACHINE™ T7 ULTRA Transcription Kit (Invitrogen, US) or the HiScribe® T7 High Yield RNA Synthesis Kit (NEB, E2040S). RNA transcripts were purified by ethanol precipitation or with the Monarch® RNA Cleanup Kit (NEB) for protoplast applications. The size and integrity of IVT RNA products were assessed by denaturing agarose gel electrophoresis. Samples (0.5–1 µg RNA) were mixed with RNA loading dye (1X buffer containing 47.5% formamide, 0.01% SDS, 0.01% bromophenol blue, 0.5 mM EDTA), heated at 65°C, and run on 1% agarose gels prepared in 1X TBE buffer alongside an RNA size standard ladder (RiboRuler High Range RNA Ladder, Thermo Scientific).

### *In vitro* protein analysis

*In vitro* translation of the synthetic mRNA occurred in wheat germ extract from Promega GmbH (L4380) and was performed as instructed by the manual. After a 3 h incubation at 25 °C, one volume of SDS loading dye was added and the samples were denatured at 100 °C for 10 min and applied onto an SDS-PAGE. The ensuing immunoblotting was performed using anti-HA from mouse (H9658, Sigma-Aldrich) and anti-mouse IgG coupled with horseradish peroxidase from goat (A4416, Sigma-Aldrich). The “Clarity Western ECL” chemiluminescence substrate from Bio-Rad (1705061) was used for peroxidase visualization and the resulting signals were detected using the Chemi Doc chemiluminescence channel with an exposure time of 15 min.

### Protoplast isolation and transfection

Protoplasts were isolated from second leaves of a 2-week-old barley plant. Isolation was performed as described by Saur et al. (2019). Only nucleic acids (mRNA and plasmid DNA) of transfection-grade purity were used for transfection. Per transfection batch, 40,000 viable cells were mixed with a total of 20 µg nucleic acids. One volume of transfection solution was added and homogenized. After a 10 min incubation at 23 °C, two volumes of wash solution were homogenized. The cells were pelleted and resuspended in an appropriate amount of regeneration buffer and incubated overnight at 25 °C. Buffer components as listed in Saur et al. (2019).

### Detached leaves mRNA spray application assay

GFP mRNA was diluted to a final concentration of 2 ng/µl in 500 µl of water and Tris-EDTA (TE) buffer. The total amount of Silwet (Silwet L-77, Momentive, US) used for each spray application was 0.2%. TE buffer diluted in 500 µl of water was used as a negative control. Up to 8 second leaves from 2-week-old barley or wheat seedlings have been harvested and placed with its adaxial side facing upwards on a square petri dish with 0,7% (w/v) water-agar to prevent them from drying out. The upper half of the detached leaves were sprayed homogeneously using flask dispensers and airbrush pistols. After spraying, the Petri dishes were closed, exposed to light and stored at room temperature. Imaging was performed using a confocal laser scanning microscope (LSM880, Carl Zeiss GmbH, Jena, Germany) 48 and 72 hours after spray application.

### CLSM sample preparation and imaging

Cross sections of barley (*Hordeum vulgare*) and wheat (*Triticum aestivum*) plants were done by hand with a razor blade and placed on an objective slide. The cross-section, as well as the adaxial and abaxial side of each cut were imaged using an LSM 880 microscope (Zeiss Microscopy GmbH, Germany) with the 488 nm laser line of an argon multiline laser (11.5 mW). Images were taken with a 40x objective (Plan-Apochromat 40_/0.8) 48 and 72 hours after spray application. Lambda stacks were created using the 32 channel GaAsP detector followed by linear unmixing with the ZEN software. Z-stacks were taken from specific areas of the sample and maximum intensity projections were produced with the ZEN software.

### Ribosome Profiling

Polysome profiling was performed following the protocol by Dong et al. (2023) [DOI: 10.1016/j.celrep.2023.112892]. Miltenyi StemMACS™ eGFP mRNA was used at a final concentration of 100 ng/µl for spraying. Samples were collected 48 hours post spraying. Total RNA from polysome fractions was reverse-transcribed using the RevertAid H Minus First Strand cDNA Synthesis Kit (Thermo Fisher Scientific). Quantitative PCR (qPCR) was performed with Luna Universal qPCR Master Mix (NEB, www.neb-online.de). The following primers were used for qPCR analysis: GFP: forward 5’-CAAGATCCGCCACAACATCG-3’, reverse 5’-GACTGGGTGCTCAGGTAGTG-3’; ACT2: forward 5’-TGACAGCCAGGTCACGTTAT-3’, reverse 5’-AGACAATTCAAAGCGGAGAGGA-3’; UBQ10: forward 5’-GGCCTTGTATAATCCCTGATGAATAAG-3’, reverse 5’-AAAGAGATAACAGGAACGGAAACATAGT-3’

## Acknowledgments

We thank Ivan Manzini and Thomas Offner from the Animal Physiology Department, University of Giessen, for their support with two-photon microscopy. We also gratefully acknowledge the valuable assistance of our employee Ann-Kathrin Hinrichs in conducting experiments.

